# Sex-Specific Stress Outcomes Following Acute and Chronic Social Defeat Stress

**DOI:** 10.64898/2026.09.25.754264

**Authors:** Erika Stewart, Alessia Manganaro, Kristin Anderson, Allie Lipshutz, Shwatina Jagnarine, Domingo Fortuna, Lillian Bryan, Labiba Aziz, Peter Rogu, Victoria Saltz, Dani Dumitriu

## Abstract

Stressful experiences are a major risk factor for the development of neuropsychiatric disorders in vulnerable individuals. Identifying features that can distinguish stress-susceptible from resilient individuals remains a central goal in stress neurobiology. Due to the routine exclusion of female subjects from stress research or the use of male-defined criteria in behavioral characterization of subjects post-stress, female stress responses and vulnerability profiles remain poorly understood. This work aimed to gain a better understanding of sex-specific stress responses to social defeat stress. Here, we used a novel paired male–female social defeat paradigm and tracked subjects longitudinally in a “two-hit” design to evaluate responses to both acute social defeat stress (ASDS) and chronic social defeat stress (CSDS), and to determine how prior stress exposure influences subsequent vulnerability. We found that both ASDS and CSDS induces changes in social avoidance/approach behavior in males but not females, as measured by a classic social interaction ratio. Prior ASDS exposure enhanced susceptibility to subsequent CSDS most clearly in males, with females displaying specific changes in corner zone occupancy. Furthermore, we observed divergent anxiety profiles between males and females in a post-stress behavioral battery. Finally, females exhibited specific stress-related locomotor responses following ASDS. Together, these findings suggest that females may adopt distinct behavioral strategies following social defeat stress.

## Introduction

While stress is a pervasive experience, individual responses to stressful events and their effect on the brain are far from homogenous. In a subset of vulnerable individuals, stressful events can trigger the emergence of neuropsychiatric disorders such as anxiety and depression while others remain behaviorally and emotionally stable following stress (Charney, 2003; Kendler et al., 1999; Kessler, 1997). Adding to this heterogeneity, there is evidence of sex differences in the response to stress and the symptomatology of such disorders (Bangasser & Cuarenta, 2021; Olff, 2017; Seney & Sibille, 2014). With mental health disorders representing a huge portion of disease burdens worldwide and decades of treatment-focused research efforts resulting in only marginally effective treatments, there is a critical need for research that is focused on both better characterizing the neurobiology of stress response phenotypes (Arias et al., 2022). More importantly, research should focus on identifying susceptible individuals prior to the onset of mental affect, to facilitate preventative interventions and promote resiliency.

CSDS is a well-validated pre-clinical rodent model of psychosocial stress that recapitulates many of the behavioral effects reported in humans, such as social withdrawal or anhedonia, that are characteristic of neuropsychiatric disturbances (Golden et al., 2011; Krishnan et al., 2007). In this model, mice experience repeated bouts of social defeat over 10 days in the form of physical attacks and other non-physical aggressive signals from a larger aggressive conspecific (Golden et al., 2011). In addition to face validity, this model effectively captures the heterogeneity of stress responses. Cohorts of genetically identical mice can be reliably stratified into susceptible and resilient subpopulations based on their behavior along a spectrum of social approach and avoidance behavior post-stress. This has enabled characterization of specific changes in susceptible versus resilience mice post-chronic stress (Christoffel et al., 2011).

It is well established that chronic stress can have long-term effects on mental health and behavior, yet stressful experiences often occur as specific isolated events. Acute stressors result in temporary increased vigilance and anxiety but do not generally lead to any long-term behavioral or psychological disruptions (Grossman et al., 2022; McCall et al., 2015). However, acute stress exposure has been shown to increases susceptibility in male mice exposed to subsequent chronic stress, a phenomenon known as stress priming or stress sensitization (Grossman et al., 2022; Weber et al., 2019). As with any salient experience, stress experiences result in memory formation, and this memory information will subsequently guide future behavior. Indeed, recent evidence has shown that disrupting memory consolidation through propranolol administration reduces stress priming (Manganaro et al., 2026). Acute stress may hold key information on an individual’s response and vulnerability to later stressors thus characterizing these early changes may be fundamental for identifying susceptible individuals prior to the onset of mental affect. The development of an acute social defeat paradigm has been crucial for characterizing early stress responses. Work from our group developed a model of ASDS where mice experience one session of social defeat over 15 minutes, designed to assess the effect of an acute stressor on behavior and brain circuitry (Grossman et al., 2022). Importantly, this model recapitulated the divergent stress responses observed following CSDS. This permits the characterization of initial responses to an acute stress experience, allowing us to track stress responses longitudinally and correlate initial stress responses with post-chronic stress phenotypes and accompanying neuropsychiatric symptomology.

A major unresolved issue in the stress field is how these responses differ between males and females. Clinical studies have consistently shown that sex is a critical variable in stress-related psychopathology, with well-documented sex differences in both the incidence and symptomatic presentation of anxiety- and depression-related disorders (Kornstein et al., 2000; McLean et al., 2011; Seney & Sibille, 2014). Despite this, preclinical rodent stress research has historically focused predominantly on male cohorts, leaving a substantial gap in our understanding of female stress responses (Merone et al., 2022; Rechlin et al., 2022). While there has been an increase in the inclusion of female subjects, the field still relies heavily on classical outcome measures originally established in male cohorts. As a result, female stress effects may be underestimated or mischaracterized if they are assessed only through male-derived behavioral criteria (Woitowich et al., 2020).

In the present study, we investigated sex-specific behavioral responses to both acute and chronic social defeat stress. First, we validated a modified version of our previously established ASDS paradigm in pairs of male and female mice using classic social approach/avoidance measures. This novel paired male-female model combines elements of previously developed models both of physical defeat and witness defeat models (Golden et al., 2011; Grossman et al., 2022; Harris et al., 2018; Iñiguez et al., 2018; Yohn et al., 2019). We then employed a longitudinal two-hit design to test whether acute stress primes males and females differently for later susceptibility to CSDS. Finally, we examined specific locomotor measures alongside an extended behavioral battery to determine whether female stress phenotypes may be better captured by behavioral features beyond the classical male-defined outcomes. Together, this approach was designed not only to characterize stress susceptibility across time, but also to determine whether sex-specific behavioral strategies shape how stress phenotypes are expressed and detected.

## Materials and Methods

### Animals

Experimental C57BL/6J 7-week-old male and female mice were received from Jackson Laboratories and group housed by sex with five mice per cage. All cages were maintained on a 12-hour light/dark cycle (light on 7am-7pm), with food and water *ad libitum*, except during the Sucrose Preference Test. All behavioral experiments began when mice were 8-10 weeks old and occurred during light hours. Retired breeder CD-1 male mice from Charles River Laboratories were used for social defeat experiments. CD-1 mice were housed in pairs separated by a metal cage divider and screened for their level aggression prior to the beginning of experiments (see below for details on aggression screening). Those classified as aggressive were used for defeat sessions and for pair-housing during ASDS and CSDS, whereas nonaggressive CD-1 mice were used for Social Interaction (SI) testing.

All experiments were conducted in compliance with National Institutes of Health Guidelines for Care and Use of Experimental Animals and approved by the Institutional Animal Care and Use Committees at Columbia University Irving Medical Center and the New York State Psychiatric Institute.

### Social Defeat Stress

#### Screening for aggression in male CD-1 mice

The level of aggression shown by CD-1 mice can vary greatly and as such CD-1 mice are screened for their level of aggression prior social defeat stress. Given that females are used in this study, we adapted the screening protocol described by Golden et al. (Golden et al., 2011), so that each CD-1 mouse is screened for aggression towards a novel C57BL/6J male and female mice for 5 consecutive days to ensure the desired level of repeated aggression is maintained over the entire duration of CSDS. Briefly, urine is collected from a different CD-1 mouse from the tested one, and 20µl of urine are placed on the urogenital regions of female and male screener mice. These male and female screener mice are then immediately added to the tested CD-1 home cage. Latency to attack and number of attacks are recorded. CD-1 mice that reach criteria for aggressive behavior are used for social defeat, whereas non-aggressive mice are kept as target mice for the SI test.

#### Acute social defeat stress

Here we used a modified version of our novel ASDS paradigm described in Grossman et al. (Grossman et al., 2022) to include male/female pairs. One hour before beginning ASDS, urine was collected from non-aggressive CD-1 mice. Twenty µL of this urine was place on the urogenital region of both male and female C57BL/6J mice and the mice were placed simultaneously into the home cage of a CD-1 aggressor for 5 minutes. This was repeated three times sequentially with no rest periods. Control male/female pairs were allowed to interact in their home cage for the duration of the defeat session and in one cohort of controls were allowed to interact with a novel conspecific instead of their defeat partner. As previously reported, (Grossman et. al., 2020) all mice were singly housed following defeat and tested for SI 60 minutes following the onset of ASDS.

#### Chronic Social Defeat Stress

We performed a modified version of the classic CSDS protocol to include male/female pairs (Berton, et al., 2006; Krishnan, et. al., 2007; LaPlant, et. al., 2010; Vialou, et al., 2010; Golden, et al., 2011). Our protocol combined elements from two other female CSDS paradigms (Harris et al., 2018; Yohn et al., 2019). In brief, experimental C57BL/6J mice were placed into the cage of a novel aggressive CD-1 mouse for 5 min daily for ten consecutive days. For the remainder of each 24-hour period male/female pairs are separated and paired-housed with a novel aggressive CD-1. A clear perforated barrier was placed between the C57BL/6J mouse and resident aggressor to allow sensory, but not physical interaction. Control C57BL/6J mice were allowed to interact in male/female pairs for 5 minutes, as in ASDS, and then pair-housed with a clear, perforated barrier. One cohort of controls were handled by the experimenter for 2 minutes. Before each daily defeat session, fresh non-aggressor CD-1 urine was applied to experimental mice as described above. At the end of 10 days, C57BL/6J mice were group-housed with prior cage mates for 24 hours. Each group-housed cage of experimental mice contained either four defeated or four control mice to avoid transfer of stress phenotype. Cages with defeated mice each contained mice from a mix of experimental groups. 24-hours following the conclusion of CSDS, experimental mice were tested for SI.

### Social Interaction Test

SI testing was performed as previously described (Golden et al., 2011). SI performed was identically following ASDS and CSDS at 1 hour and 24 hours, respectively. Each single experimental mouse was placed into an open-field arena (45 cm L x 45 cm W x 30 cm H) with a removable wire-mesh enclosure in clear Plexiglas (10 cm L x 6.5 cm W x 42 cm H) placed against one of the inner walls. Mice were allowed to explore and habituate to the arena for 150 seconds without a target mouse in the enclosure (No Target session). After that, experimental mice were removed and a novel non-aggressive CD-1 “target” mouse was placed in the wire-mesh enclosure. The experimental C57BL/6J mouse was reintroduced to the arena for another 150 seconds (Target session). Open-field arenas and wire-mesh enclosures were wiped with 70% ethanol solution between sessions. The test was conducted during the light phase (between 13:00 and 19:00) under infrared light conditions. The total time spent in the “interaction zone” (IZ) defined as a 24cm x 8cm corridor surrounding the wire-mesh enclosure, and in the “corner zone” (CZ) defined as the cumulative area of the two corners (9cm x 9cm each) opposite to the interaction zone, was calculated. In addition, we calculated the SI and corner ratios as the absolute time spent in the respective zone when target is present divided by the absolute time spent in the zone when target is absent (Grossman, et. al. 2020).

### Behavioral Battery

#### Light-Dark Box Test

Experimental mice were placed into an open-field arena (42 cm L x 42 cm W x 38 cm H) consisting of two connected compartments (Smart Field Open Field System, Kinder Scientific), one brightly illuminated light-zone and one enclosed and dark-zone (42 cm L x 21 cm W x 30 cm H). Mice could freely move between zones through a doorway (9cm W x 11cm H). Mice were placed in the light zone first and allowed to freely explore both compartments for 10 min. IR beam sensors located along the X and Y axes automatically captured animal movements between compartments (Motor Monitor Software, Kinder Scientific). Behavior was also recorded using an overhead camera. The time spent in the light and dark zone and latency to enter was quantified.

#### Three-Chamber Sociability Test

We evaluated sociability in mice using the three-chamber social test (Moy et al., 2004). Experimental mice were placed in the center of a three-chamber apparatus (42cm W x 63cm L x 30 cm H) and allowed to habituate for 5 minutes. Each of the two outer chambers (42cm W x 21 cm L) contained a cylindrical interactor (9cm W x 28cm H) with either a novel, non-experimental C57BL/6J mouse of the same sex (social chamber) or a novel object (non-social chamber). The partitions between each chamber were opened and the experimental mouse was allowed to freely explore the whole apparatus for 10 minutes. The time spent in each chamber was recorded. The sociability index was calculated as time spent in the social chamber minus time spent in the non-social chamber over total time in either zone. A sociability index of zero indicates no preference between the social and non-social cues. A positive index reflects a preference for the social cue, while a negative index suggests a preference for the non-social cue.

#### Elevated Plus Maze

We conducted the elevated plus maze as described by (Kraeuter et al., 2019). C57BL/6J mice were placed in the center of a plus maze apparatus (88cm W x 22cm L x 50cm x 0.5 H) made from black Plexiglas with two open arms (35cm L x 5cm W) and two closed arms (35cm L x 5cm W x 15 cm H) and an intermediate middle zone between arms (9cm W x 9cm L x 0.5 cm H). Mice were allowed to explore freely for five minutes, and their movements were video recorded.

#### Sucrose Preference Test

For the duration of sucrose preference testing, experimental mice were provided with food *ad libitum* and were singly housed to allow for quantifiable fluid consumption measurements. Mice did not have access to water except for the two standard water bottles placed inside each cage. On day one, each water bottle contained only water in order to establish any baseline preference over the course of 24 hours. On day two, one of the two bottles was filled with 1% sucrose water solution and the initial weight of both bottles noted. After 24-hour period, the weight of each bottle was measured to calculate total fluid consumption, and mice were returned to their home cages with their previous cage mates.

#### Composite z-anxiety score

Each anxiety behavioral measurement (Time spent in light zone, Time spent in open arms and Sociability Index) was first converted into a z-score relative to the appropriate control group, with the formula Z-score = (X – mean of controls / SD controls). The resulting standardized values were then averaged to generate a composite anxiety-like z-score. Positive z-scores indicated greater anxiety-related behavior and negative scores indicated less anxiety-related behavior relative to the reference control group.

Only animals that had data for all three of these behaviors were included in the composite score, as some animals were missing data for one or more measures.

### Locomotor analysis

Following accurate video analysis, individual animal position during the SI test (25Hz, 40ms sampling interval) was extracted in timestamped X-Y coordinates using Ethovision XT (Noldus Information Technology). For each frame, instantaneous speed was computed as the Euclidean displacement between consecutive XY centroid positions divided by the inter-frame interval: speed = √ (Δx² + Δy²) / Δt (cm/s). Instantaneous acceleration was defined as the absolute rate of change in speed between consecutive frames: acceleration = |speed_t − speed_{t−1} | / Δt (cm/s²). For each animal and session, the mean value of each kinematic variable was computed across all valid tracking frames and used as the unit of analysis. Group differences were assessed by two-way ANOVA with Sex (male, female) and Condition (control, defeat) as between-subject factors, followed by Tukey-corrected pairwise comparisons.

Delta values for speed and acceleration were calculated as the change in these values between the No Target and Target trials for each subject Delta Δ = (No Target Speed – Target Speed)

### Experimental design, timeline, groups and sample sizes

This investigation aimed to examine the effects of ASDS in male-female pairs, in addition to examining the priming effect of ASDS for increased susceptibility to CSDS in the same mouse pairs (Fig 1B). To address these questions, we designed a multi-stage experiment that included both ASDS and CSDS followed by SI testing (Fig 1A). In addition, two extensive post-stress behavioral batteries were included to examine long-term behavioral alterations resulting from acute and chronic stress (Fig 1A). Animals were randomly assigned to one of seven primary groups that completed all or selected stages of the experimental protocol below (Fig 1C).

In the first stage, two groups underwent either ASDS (ASDS n = 57 males, 50 females) or a non-stressful control social interaction (ASDS Control n = 26 males, 23 females), followed by an SI test one hour later. One week later, some of those mice underwent a behavioral battery consisting of the light/dark box (L/D), three-chamber social test (3C), elevated plus maze (EPM), and sucrose preference test (SPT) (ASDS Control n=9 males n=9 females, ASDS n = 15 males, n=15 females). At this stage, a group of mice completely naïve to ASDS was added to also complete the behavioral battery (BEH + CSDS n = 9 males, 9 females). One week following the battery, these groups (ASDS Control n = 9 males, 9 females, ASDS + CSDS n= 15 males, 15 females, BEH +CSDS n = 9 males, 9 females) underwent CSDS. Two further groups of previously untested mice were introduced at this stage; one group went to experience CSDS (CSDS n = 23 males, 23 females) and one served as control group (Control CSDS n = 25 males, 25 females) to social defeat stress. One-week post-CSDS, some animals underwent the second behavioral battery (Control n=10-15 males, 8-10 females, CSDS n = 20-30 males, 20-30 females, ASDS + CSDS n=10-15 males, 8-11 females). For evaluation of stress priming, all mice that underwent CSDS but that did not have a prior ASDS experience (ASDS control, Battery + CSDS, and CSDS, n=41 males, 41 females) were combined into one CSDS group, after comparison analysis between the groups excluded significant differences in the SI and corner ratio (Fig S2).

In literature, social defeat studies often differ in the control conditions used. In some reports, control mice interact with novel same-sex conspecific mice for the duration of a defeat session (Grossman et al., 2022) or some interact with non-aggressive CD-1 (Takahashi et al., 2017). In other cases, control mice are handled by the experimenter (Schuler et al., 2025; van Doeselaar et al., 2021). Control housing conditions also varies with some control mice being singly house while others are cohoused with a same-sex conspecific (Harris et al., 2018; Takahashi et al., 2017). Our primary control condition consisted mainly of male/female pairs interacting in their home cage for the duration of the defeat session (Yohn et al., 2019). However, we took into considerations other well-used control conditions. In a separate cohort CSDS controls (n = 10 males, 10 females) were handled daily for 2 minutes by the experimenter and returned to their home cage, co-housed with a same-sex conspecific. The CSDS condition (n = 11 males, 11 females) for this cohort was instead unchanged.

For the ASDS, one cohort of mice had different control conditions (controls: n= 17 males, 14 females, defeated: n = 42 males, 35 females), with mice undergoing 3 x 2 min interactions with a novel conspecific. This cohort was combined with the main dataset as a larger sample size was required for some of our analysis but are also presented in isolation in supplementary (Fig S6). In each case, the same overall differences between control and defeat groups were observed, supporting the consistency of our findings across independent cohorts of mice with various control conditions (Fig S6).

**Figure 1:**
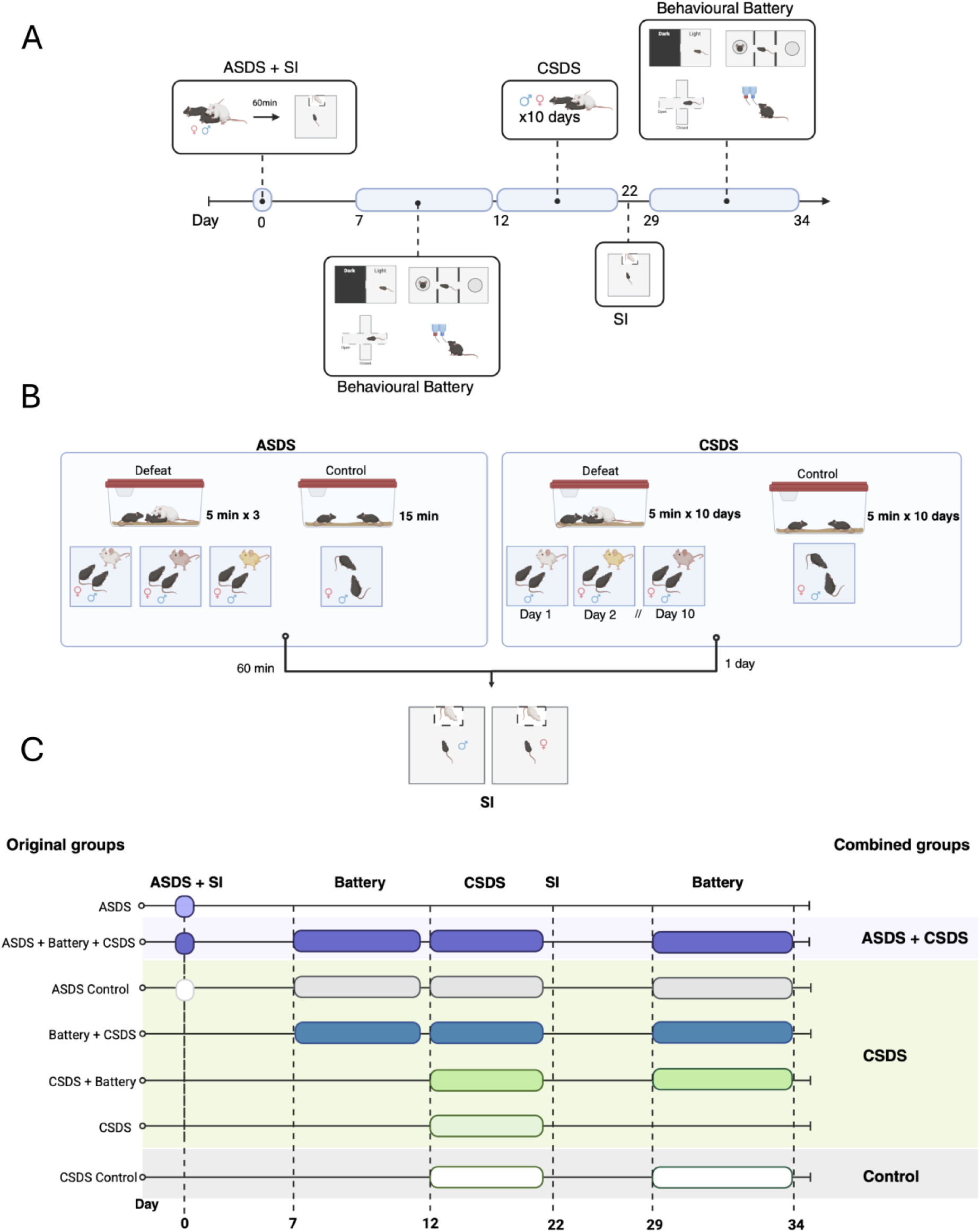
Experimental design, timeline and groupings. (A) Experimental design and timeline. (B) Schematic depiction of ASDS and CSDS protocol (C) Schematic of experimental groups. Schematics created in BioRender.

### Analysis and statistics

A video-tracking system (Ethovision 18X.0, Noldus Information Technology) was used to record the movements and behavior during each behavioral test.

Data analysis and statistics were conducted using GraphPad Prism 11 (GraphPad software). Group means (Control v Defeat and Male vs Female) were analyzed using two-way ANOVA followed by Bonferroni or Tukey adjusted post hoc comparisons, as appropriate. Within-group comparisons across experimental time points (ASDS SI vs CSDS SI) or session conditions (no target vs target) were analyzed using repeated-measures ANOVA or paired *t*-tests, as appropriate. Effect sizes for group comparisons were quantified using Hedges’ g. Associations between continuous variables were assessed using Pearson’s correlation and differences in categorical proportions were assessed using Fisher’s exact test. All data were graphed as mean + standard error of the mean (SEM). An alpha level of 0.05 was used as a criterion for statistical significance.

Schematic designs created using BioRender

## Results

### ASDS induces social avoidance in male but not female mice

Our first objective was to validate the use of our previously developed novel ASDS paradigm in pairs of male and female mice. Combining a few components from existing female defeat models, we paired male and female C57BL/6J mice and subjected them to three consecutive five-minute defeat session, with three different resident aggressive CD-1 mice. Immediately prior to placing the intruder pairs into the CD-1 cage, both mice had twenty µl of CD-1 urine applied to their urogenital region. One-hour post-stress mice underwent an SI test with a non-aggressive CD-1 mouse to measure social approach and avoidance behavior (Fig 2A). To test whether ASDS induced social avoidance in male and female mice, we first evaluated the SI ratios between control and defeated groups (Fig 2C). Using a two way ANOVA we found an overall significant main effect of both sex (F (1,150) =20.06, p<0.0001; Fig 2C) and stress (F(1,150)=4.681, p=0.0321; Fig 2C) in addition to a significant interaction between stress and sex (F (1,150)=7.738, p=0.006; Fig 2C). Consistent with our previous findings (Grossman et al., 2022), the multiple comparison analysis revealed a significant decrease of the SI ratios in ASDS males compared to controls (p=0.0026; Fig 2C). The same effect was not present in female mice (p>0.9999; Fig 2C), where control and defeated mice were undistinguishable. Importantly, female controls had significantly lower SI ratios than male controls (p=0.0001, Fig 2C). Furthermore, these results are reflected in the analysis of the absolute time the animals spent in the IZ without and with the CD-1 target. A two-way repeated measures ANOVA (RMANOVA) found a significant interaction between stress and session (no target vs target) in the absolute time males spent in the IZ (F (1,77)=13.00, p= 0.0006; Fig 2D), as well as a significant main effect of session (F (1,77) =25.98, p<0.0001, Fig 2D) and stress (1,77)=4.625, p=0.0346;Fig2D). The multiple comparisons analysis indicated that controls spent significantly more time in this zone in presence of the CD-1, indicating natural social approach behavior (p<0.0001; Fig 2D), whereas ASDS male spent significantly less time than controls in the zone during the target session (p=0.0002;Fig 2D). For females, we did not find any significant change on the time spent in the IZ between sessions for either control or ASDS groups (Fig 2E), suggesting that even in absence of stress, female mice did not exhibit an approach behavior. We then proceed to examine the time the animals spent in the CZ opposite to the interactors, as a possible behavioral sign of avoidant and anxious behavior. The two-way ANOVA identified a significant main effect of sex on the corner ratios (F (1, 150) =4.365, p =0.0384; Fig 2F), although post-hoc comparisons did not identify a significant difference between groups in either sex. Instead, the analysis on the time spent in the CZ found a significant interaction between stress and session in males (F(1,77)= 6.945, p = 0.0102; Fig 2G) and post-hoc analysis found significant difference between controls and ASDS mice during the target session (p=0.0343;Fig 2G). In female we observed identified a main effect of session (F(1,71) = 8401, p=0.0050;Fig 2H) where ASDS females showed a significant increase in time spent in corner zones between session (p=0.0219;Fig 2H).

As previously reported, defeated male mice show a large degree of variability in their behavior and can be further stratified into susceptible and resilient subgroups after ASDS (Grossman et al., 2022). Control mice generally display social preference and greater approach behavior illustrated by an SI ratio > 1. Therefore, defeated subjects with an SI ratio ≥ 1 are defined as resilient and < 1 are deemed susceptible. We plot the distribution of SI and corner ratios for control and ASDS groups (Fig 2I-L). We stratified males into resilient and susceptible sub-groups based on this classification however female mice were not further stratified as they did not demonstrate a significant difference in SI compared to controls (Fig 2I-L). Notably, we observe a large proportion of female control mice that fall below 1 indicating that this stratification criteria are not appropriate for female mice as controls did not demonstrate social preference (Fig 2J).

7 days post-ASDS mice underwent an extensive behavioral battery to examine the effect of ASDS on long-term anxiety, sociability and depression-like behaviors (Fig 2M). ASDS had no significant effect on behavior in the L/D, 3C, EPM or SPT in either males or females (Fig S1A-D). Since multiple anxiety-related assays were included in this behavioral battery, we calculated an overall composite score to capture distributed behavioral effects and provide a broader index of the stress-related phenotype that may not be evident from individual measures alone (Guilloux et al., 2011). Each behavioral measure was first converted into a z-score relative to the appropriate control group, and the resulting standardized values were then averaged to generate a composite anxiety-like z-score. No significant difference between control and stress groups was identified, however a significant difference was observed between male and female stress groups (p= 0.0073; Fig 2N) indicating that the combined behavioral profile differed between males and females, where males demonstrated lower values indicating increased anxiety and females demonstrated higher values indicating less anxiety.

These findings highlight ASDS induces obvious changes in social approach/avoidance behavior in males but not females.

**Figure 2:**
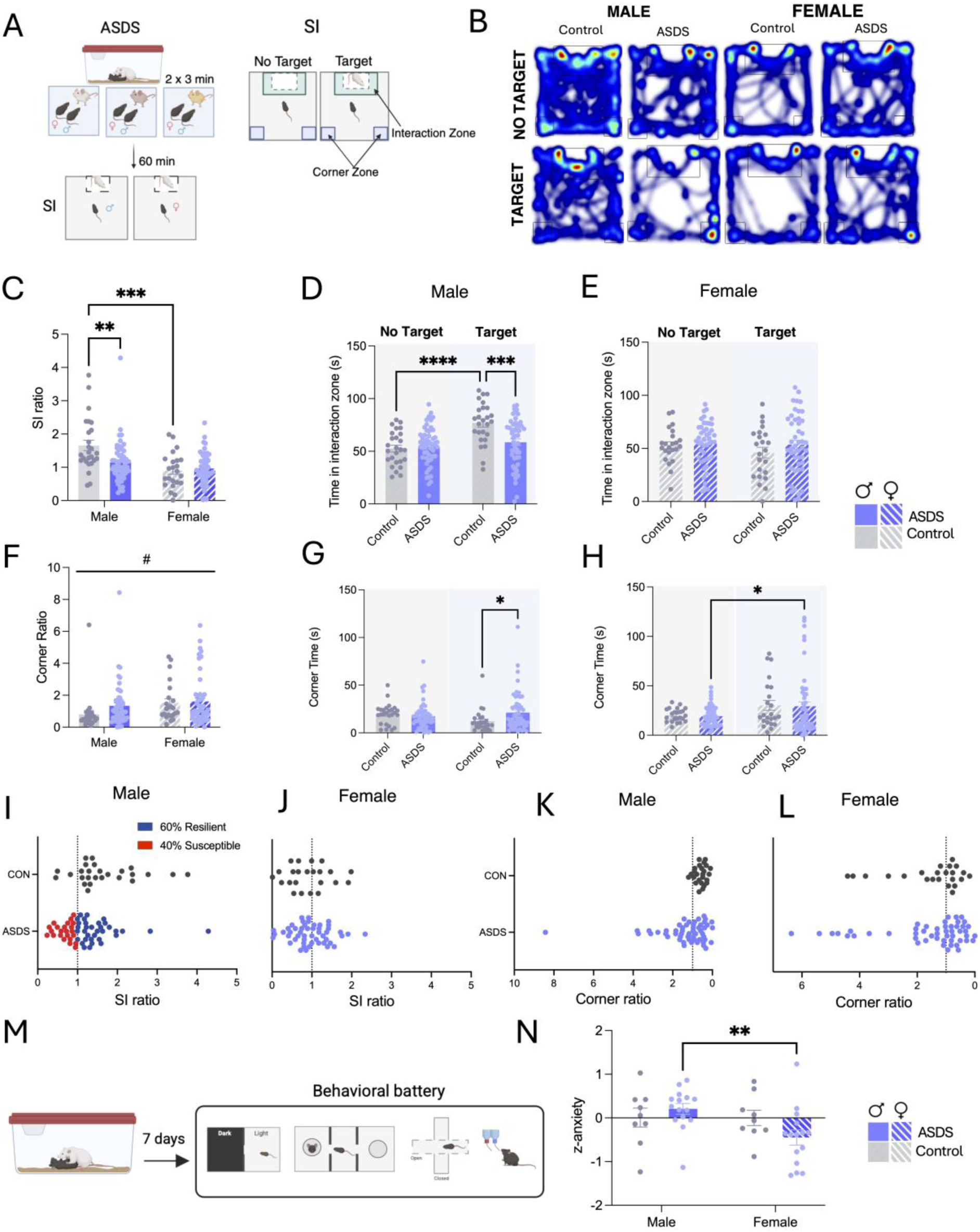
ASDS induces social avoidance in male but not female mice. (A) Schematic of the ASDS and SI workflow. (B) Representative heatmaps of cumulative occupancy in SI arena across essions. (C) Quantification of SI ratio in control and defeated mice. (D-E) Quantification of raw time pent in the IZ across sessions in either male or female control and defeated mice. (F) Quantification f corner ratio in control or defeat mice. (G-H) Quantification of raw time spend in CZ during target and no target trials in either male or female control and defeated mice. (I-J) Distribution of SI ratio in male and female mice and the proportion of each stress phenotype (K-L) Distribution of corner ratio n male and female mice. (M) Schematic of the behavioral battery 1-week post ASDS (N) Quantification of composite anxiety score in control and defeated mice. Data presented as mean +/-SEM. Statistical comparison performed using (C,F,N) Two-way ANOVA with Bonferroni post-hoc D,E,G,H) Two-way RMANOVA with Bonferroni post-hoc. Main effect: #P < 0.05, Post-hoc: *P < 0.05, *P < 0.01, *** P < 0.001, **** P < 0.0001.

### ASDS primes for increased susceptibility following CSDS but drives a sex-specific behavioral phenotype

While an acute stress experience alone does not give rise to long-term behavioral alterations, we previously reported that exposure to ASDS can increase sensitivity to subsequent stress resulting in increased susceptibility following a second exposure (Grossman et al., 2022; Manganaro et al., 2026). Our next objective was to examine the effect of chronic stress in our paired male-female model, in addition the effect of stress priming on both males and females. We first examined the effect of CSDS on mice grouped by their prior experience (Fig S2A). Two-way ANOVA detected a main effect of stress group (F(4,152)=8.018, p=<0.0001;Fig S2B) and sex F(1,152)=6.399,p=0.0124;Fig S2B) on SI ratio. Multiple comparisons revealed significant differences between CSDS control and ASDS Control (p=0.0009), ASDS +CSDS (p=<0.0001) and Battery + CSDS (p=0.0337) groups (Fig S2). We also found a significant difference between CSDS and ASDS + CSDS groups (p=0.0413;FigS2). We found no significant difference between groups that underwent CSDS with a prior behavioral battery and/or a control social interaction but did not have a prior ASDS experience and therefore we combined these into one CSDS group for further analysis (Fig 3). To test this end, we compared control mice that experienced no stress with mice that experienced CSDS, or mice that had both ASDS followed by CSDS (Fig 3A). Using a two-way ANOVA to evaluate the SI ratio between groups we found a main effect of both stress (F (2, 155) =11.67, p=0.000; Fig 3B) and sex (F (1, 155)=8.457, p=0.0042; Fig 3B) in addition to a significant interaction (F (2, 155) = 4.547, p=0.0120; Fig 3B). Multiple comparison analysis revealed that in males, both CSDS and ASDS + CSDS showed significant decrease in social approach as illustrated by a significant decrease SI ratio compared to control mice (p=0.003,p=<0.0001 respectively; Fig 3B). In contrast, we did not find any difference between female stress groups, but consistent with what we observed following ASDS, we found a significant difference in SI ratio between control male and females (p = 0.0001;Fig3B). In males, while both stress groups illustrate a significant decrease in SI ratio, the effect size was larger in the ASDS + CSDS (Hedges g = 1.43) compared to CSDS (Hedges g = 0.86; Fig 3B), coherent with a priming effect of ASDS. While we did not observe any difference in SI ratio between stress groups in female mice, we found a main effect of stress on corner ratio (F(2,155)= 9.349,p=0.0001;Fig 3E) where female ASDS + CSDS mice had a higher corner ratio compared to both controls (p=0.0333) and CSDS (p=0.0426) females (Fig 3E). In males, we found a significant difference in corner ratio only between controls and ASDS + CSDS (p=0.0177; Fig 3E). These differences in SI and corner ratio are also reflected in the absolute time spent in the IZ and CZ across sessions. In males, a two-way RMANOVA of time spent in IZ detected a significant effect of session (F(1,78)=4.354, p=0.0402;Fig 3C) and an interaction between session and stress (F(2,78)=8.89, p=0.0003;Fig 3C). Multiple comparison analysis revealed that control males significantly increased their time spent in IZ zone between sessions (p=<0.0001;Fig 3C). During the target session, control males also spend significantly more time in IZ compared to ASDS + CSDS mice (p=0.0125;Fig 3C). In females, there was a significant main effect of session (F(1,78) = 4.52, p=0.0367; Fig 3D). Female ASDS + CSDS mice spent significantly less time in the IZ during the target trial compared to the no-target trial (p=0.032;Fig 3D). Two-way RMANOVA of raw time spent in the CZ found a significant interaction between target and stress in both males (F(2,78) = 6.54, p=0.0024;Fig 3F) and females (F(2,78) = 5.57, p=0.055;Fig 3G) where stress significantly increased time spent in the CZ in both sexes during the target trial. We plot the distribution of SI and corner ratio for males and female mice (Fig 3H-K). Males were stratified into resilient and susceptible subgroups based on SI ratio threshold of 1 (Fig 3H). Females did not display a difference in SI ratio post-stress and we again observed a large proportion of controls with an SI ratio below 1 (3I). However, females did display a difference in corner ratio, therefore stratifying females based on corner ratio may better represent their stress phenotype and for all future analysis females were stratified based on corner ratio threshold of 1 (Fig 3K). Using Fisher’s exact test, we quantified the proportion of each phenotype and found a significant difference in the proportion of susceptible mice in male ASDS + CSDS group compared to CSDS group (Odds ratio=4.91, p=0.017; Fig 3L) but females showed similar proportions in both groups (Fig 3L). The priming effect of ASDS in males is further illustrated by comparing SI ratios post-ASDS and post-CSDS demonstrating a significant decrease in SI ratios post-CSDS (t(14)=2.336,p=0.039;Fig 3M). As previously reported this ASDS paradigm has a strong positive predictive value (PPV) for susceptibility in male mice, 100% of male mice classified susceptible post-ASDS were also susceptible post-CSDS (Fig 3M) (Grossman et al., 2022). The PPV is not as strong in females and their priming effect is only marginally significant (t(14) = 2.060,p=0.0585;Fig 3N).

Seven days post-CSDS mice underwent second behavioral battery to examine the effects of CSDS on anxiety and depression-like behaviors. We found a main effect of stress on males in the time spent in the open arms of the EPM (F(2,65)=3.92, p=0.0246; Fig S3C) suggesting an increase in long-term anxiety-like behavior in male mice following CSDS. We stratified mice by stress phenotype and found a significant difference between control and both resilient and susceptible mice in EPM time in arms (p=0.045, p=0.0108 respectively; Fig S3G). We also observed a significant difference between susceptible males and females in their sociability index in the 3C (p=0.021; Fig S3F). We found no significant effect of having a prior behavioral battery on outcomes in the second battery (Fig S4).

Two-way ANOVA of composite behavioral z-score detected a main effect of sex (F(1,68)= 8.184, p=0.0056;Fig 3O) suggesting a sex-dependent divergence in the anxiety phenotype post-stress. Multiple comparison analysis revealed a significant difference in composite anxiety score between male and female CSDS (p=0.0013;Fig 3O) and ASDS + CSDS (p=0.0212;Fig 3O) groups again highlighting a sex-specific behavioral profile following stress.

The longitudinal design of this experiment enabled us to assess of whether social approach/avoidance behaviors following acute stress predicted later anxiety-related outcomes after CSDS. Pearson correlation revealed a significant positive correlation between SI post-ASDS and time spent in the light zone during the L/D test post-CSDS in male ASDS + CSDS mice (r=0.5482, p=0.034; Fig S3I-J) but not females, suggesting a relationship between SI-post ASDS and some anxiety-like behaviors post-CSDS in males.

Together these data show that females display evidence of social avoidance post-CSDS and ASDS sensitizes females to CSDS, but their behavioral profile is different to that of males.

**Figure 3:**
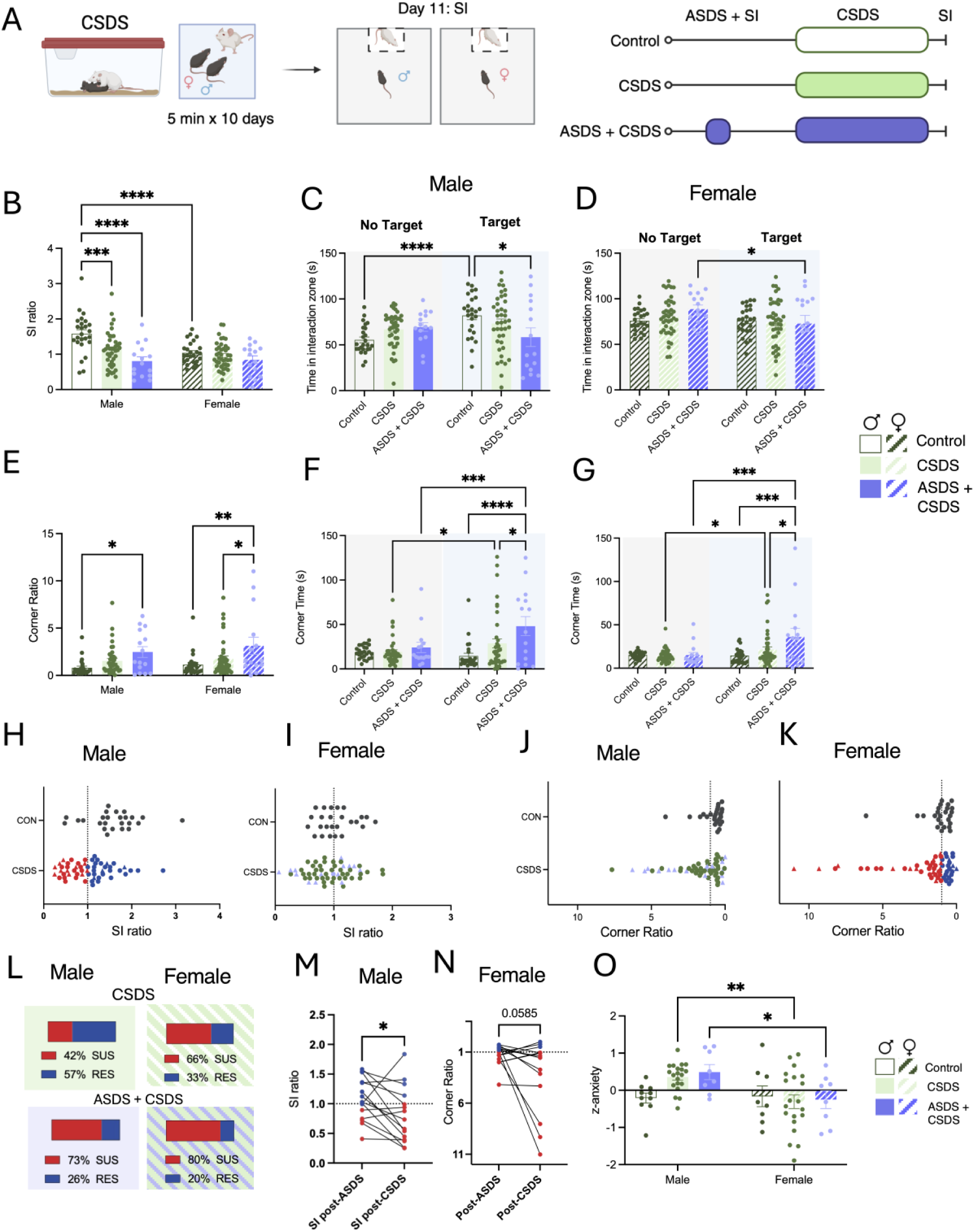
Acute stress exposure primes for increased susceptibility to chronic social defeat stress. (A) Schematic of the CSDS and SI timeline and experimental groups. (B) Quantification of SI ratio in control, defeat or double defeat mice. (C-D) Quantification of raw time spend in IZ across sessions in either male or female control, defeated or double defeated mice. (E) Quantification of corner ratio in control, defeat or double defeat mice. (F-G) Quantification of raw time spend in CZ across session in either male or female control defeat or double defeated mice. (H-I) Distribution of SI ratios in control and defeated mice (J-K) Distribution of corner ratios in control and defeated mice. Circle symbol indicates mice that underwent CSDS only and triangle symbol indicates mice that underwent both ASDS + CSDS (L) Proportion of each stress phenotype in male and female CSDS or double defeat groups. (M-N) Quantification of SI (male) or corner ratio (Female) post-ASDS and post-CSDS in the double defeat group. (O) Quantification of composite anxiety score in control, CSDS and ASDS + CSDS group. Data presented as mean +/-SEM. Statistical comparison performed using (B,E,O) Two-way ANOVA with Bonferroni post-hoc (C,D,F,G) RM Two-way ANOVA with Bonferroni post-hoc (L) Fisher’s exact test (M,N) Paired *t* test *P < 0.05, **P < 0.01, *** P < 0.001, **** P < 0.0001.

### Stress differentially alters locomotion during an SI test in male and female mice

Given that females did not consistently display a decrease in social approach behavior, we sought to evaluate other behavioral measures as candidate signatures of stress that might complement standard SI metrics. Growing evidence suggests that females may express stress- and fear-related states using different behavioral strategies than males, including changes in speed and movement patterning rather than simple reductions in social approach (Gruene et al., 2015; Schuler et al., 2025). We examined locomotor behavior (speed and acceleration) during the SI test (Fig 4A). Two-way RMANOVA revealed that all mice show reductions in speed and acceleration across sessions (Fig S5). We calculated the delta value for this change in locomotion across SI test sessions post-ASDS and a two-way ANOVA detected a main effect of sex (F(1,148) = 8.436, p=0.0042;Fig 4B) and stress (F(1,148) = 10.17, p=0.0017;Fig 4B) for delta speed and comparably a main effect of sex (F(1,148) = 15.28, p=0.0001;Fig 4C) and stress (F(1,148) = 4.797, p=0.0301;Fig 4C) for delta acceleration. Multiple comparison analysis indicated that defeated females differed from control females in both delta speed (p=0.0058;Fig 4B) and delta acceleration (p=0.0343;Fig 4C), whereas no such differences were observed in males, suggesting that ASDS elicits a female-specific adaptation in locomotor responses. These findings indicate that the female response to ASDS may be characterized by maintenance of higher locomotor speed, consistent with a distinct post-stress behavioral strategy relative to males. When stratified by stress-phenotype, both resilient and susceptible males show significantly lower speed values during the target trial compared to controls (p=0.0101 and p=0.0007, respectively; Fig 4D), but no such differences were found in females. Linear regression analysis indicated a significant relationship between corner ratio and speed in defeated females (r=-0.3464 p=0.0137;Fig4F) but not controls. In contrast, following CSDS, delta speed and acceleration did not differ between control and defeated mice in either sex, indicating that this acute female-specific effect was not maintained after chronic stress (Fig 4G-H). This interpretation was further supported by stratification according to stress phenotype where males and females showed a similar decrease of speed post-stress independent of stress phenotype. Two-way ANOVA found a main effect of stress (F(2,114) = 2.13, p<0.0001; Fig 4I) and sex (F(2,114) = 18.87, p< 0.0001; Fig 4I). Multiple comparison showed that defeated females maintained higher target-trial speed than defeated males irrespective of stress phenotype (p=0.01 mRES vs fRES, mSUS vs fSUS, p=0.0014; Fig 4I). Post-CSDS linear regression analysis showed no significant group differences in slope for the relationship between sociability ratios (SI or Corner ratio) and speed but did reveal a significant difference in elevations for both males (F(2, 56) =10.49, p=0.0001;Fig4J) and females (F(2,56)= 6.776, p=0.0023;Fig4K) suggesting group differences in overall locomotor output independent of sociability ratios.

Taken together, the overall pattern of results supports the idea that females respond differently to social defeat even when conventional social interaction ratios do not show a robust group effect.

**Figure 4.**
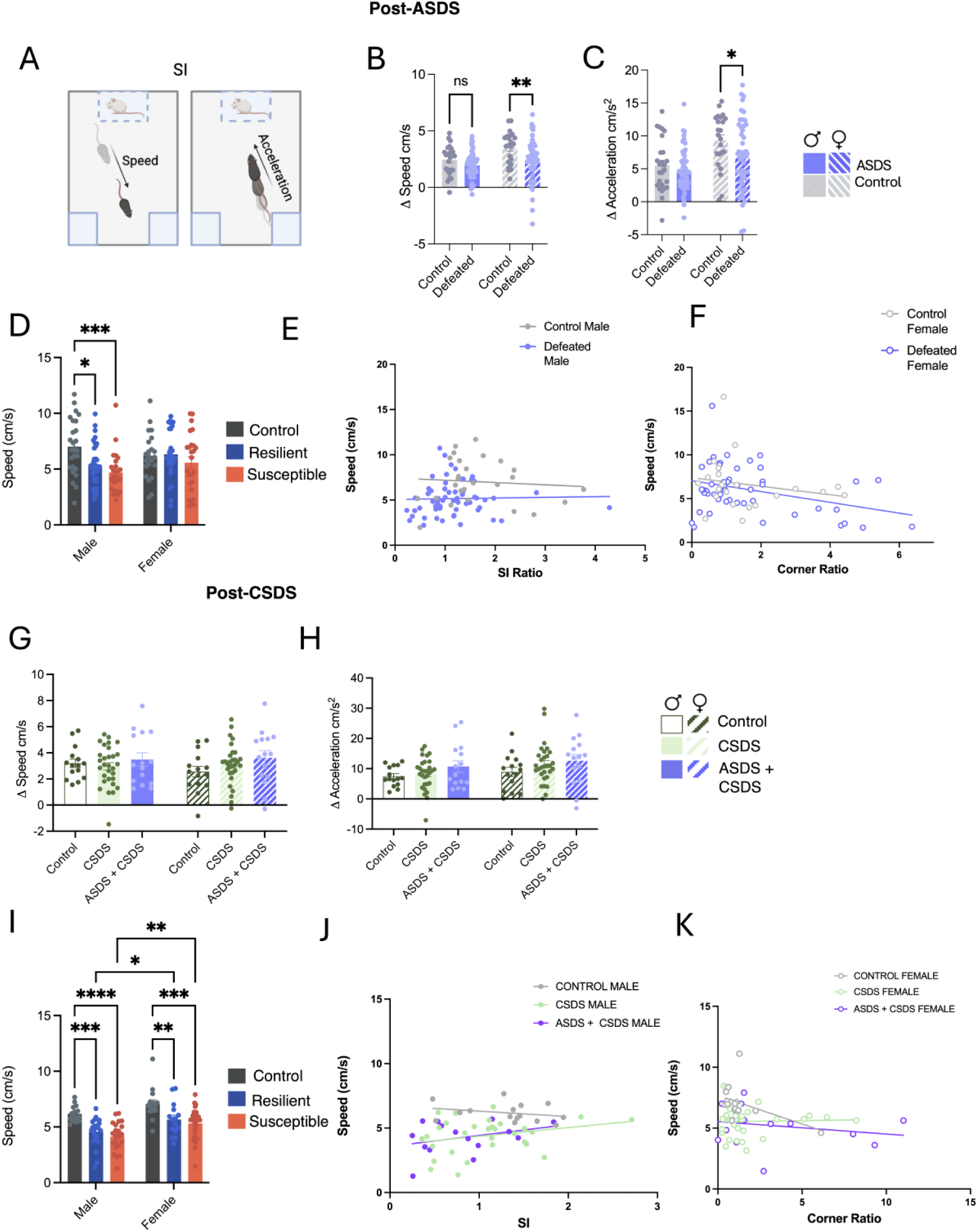
Sex-specific locomotor responses to acute and chronic social defeat stress. (A) Schematic of locomotor measures analyzed during the SI test, including speed and acceleration across no-target and target trials. (B-C) Post-ASDS change (delta Δ) in speed (B) and acceleration (C) in male and female control and defeated mice. (D) Stratification of post-ASDS target-trial speed by stress phenotype in males and females. (E-F) Linear regression of SI or corner ratio versus speed following ASDS in control and defeated male and female mice. (G-H) Post-CSDS change (delta Δ) in speed (F) and acceleration (G) in control, CSDS, and ASDS + CSDS groups in males and females. (I) Stratification of post-CSDS target-trial speed by stress phenotype in males and females. Females maintained higher target-trial speed than males across stress phenotypes. (J-K) Linear regression of SI or corner ratio versus speed following CSDS in control, CSDS, and ASDS + CSDS groups in males and females. Data are presented as mean +/-SEM. Statistical comparison performed using (B-D,G-I) Two-way ANOVA with Bonferroni post-hoc (E,F,J,K) Simple linear regression and Pearson correlation *P < 0.05, **P < 0.01, *** P < 0.001, **** P < 0.0001.

## Discussion

Within social stress research, a major limitation of established psychosocial stress models is the low level of female-directed aggression from CD-1 aggressor mice. Several alternative models have been developed to address this, including DREADD-based activation of the ventromedial hypothalamus, which induces aggression toward both male and female mice and produces comparable stress-related outcomes in females (Takahashi et al., 2017), and application of male aggressor urine, which promotes female-directed aggression (Harris et al., 2018). Witness defeat, in which mice observe another mouse undergoing social defeat, also induces behavioral and neurobiological changes similar to direct defeat and has been successfully applied to females (Schuler et al., 2025; Warren et al., 2013). More ethologically relevant approaches include using naturally aggressive strains, such as Syrian hamsters, or intact Swiss Webster females following co -housing with castrated males, which display aggression toward female intruders (McCann et al., 2019; Newman et al., 2019).

Despite these advances, a key limitation remains in that post-stress classification criteria are largely based on male cohorts and may not accurately capture stress-related changes in females. Here, we developed a paired male-female model combining elements of the urine, chronic non-discriminatory social defeat, and witness defeat models (Harris et al., 2018; Iñiguez et al., 2018; Yohn et al., 2019). Both males and females receive twenty µL of CD-1 urine before being placed together in a CD-1 home cage, resulting in robust aggressive attacks toward both sexes.

Using this model, we reproduced a robust acute social avoidance phenotype in males but not in females. Second, ASDS primed for enhanced susceptibility to later CSDS most clearly in males, consistent with the idea that a single acute stressor can lower the threshold for future maladaptive responses. Further, our composite anxiety score highlights a sex-specific anxiety profile post-stress. Finally, despite less changes in classic SI metrics, movement-based analyses captured a female specific stress response.

The lack of change in social approach/avoidance we observed in females following social defeat is coherent with other findings (van Doeselaar et al., 2021). Indeed, 21 days of CSDS produced no generalized social avoidance in female subjects (van Doeselaar et al., 2021). This highlights the limitations of relying exclusively on traditional SI-based criteria to assess female stress phenotypes. Females in this model experienced the same repeated aggressive encounters and defeat-associated sensory cues as their male partners, yet they did not consistently exhibit the same decrease in SI ratio seen in males (Fig 2C and 3B). This is bolstered by recent findings that do report a significant decrease in SI ratio in females following chronic witness defeat in a large sample size (n = 174-230), however the authors highlight that this effect was much larger in male subjects and their large sample size enabled them to capture this effect in females, stating that a more standard sample size such as the sample size in this study would not be sensitive enough to capture these differences (Schuler et al., 2025).

We performed an extensive post-stress behavioral battery to further characterize anxiety and depression like behaviors post-stress. While we found no differences in anxiety or depression like behaviors post-acute stress in any of the individual test (Fig S1), which is consistent with previous findings that acute stress does not give rise to long term behavior alterations, we did find a divergence between males and female in overall behavioral profile (Fig 2N). By integrating multiple assays, the composite score captured a sex-dependent divergence that was not always obvious from individual tests alone. In males, higher composite scores indicated a shift toward a more stress-like behavioral profile relative to male controls. In females, the composite shifted less strongly or in the opposite direction relative to female controls suggesting that the organization of the behavioral response differs by sex and that females may adopt distinct behavioral strategies after stress (Fig 2N). The observation that the composite z-anxiety score detected a group difference when individual raw measures did not reflect the distributed nature of the stress phenotype. Behavioral assays such as EPM, L/D, and sociability each capture overlapping but non-identical aspects of affective behavior, and each assay is subject to substantial within-group variability. By standardizing measures relative to controls and combining them into a summary score, the composite may provide a more sensitive index of the overall behavioral impact of stress. Post-CSDS we did observe some specific changes in anxiety-like behavior in males (Fig S3), but we also observed similar divergence in anxiety-like behavior between males and females, reinforcing a clear difference in behavioral profile (Fig 3O).

Another key observation from these results is the presence of some baseline behavioral differences between males and females, highlighting an important consideration when assessing sex differences in behavioral responses to stress. Female controls displayed lower levels of social approach compared to male controls (Fig 2C), and therefore the absence of a decrease in sociability following stress should not necessarily be interpreted as evidence that females were unaffected by stress. Similarly, we observed significant differences between control males and females in their time spent in the open arms of the EPM (Fig S2C). These findings emphasize the importance of considering baseline sex differences when interpreting stress-induced behavioral changes.

A growing body of evidence that suggests sexually divergent changes in movement in response to fear and stress stimuli, motivated us to analyze locomotion during the SI test (Gruene et al., 2015; Schuler et al., 2025). Following fear conditioning, female rats display a novel active fear responses in the form of darting behavior, which is described as rapid, high velocity movement (Gruene et al., 2015). More recent work has identified female specific maintenance of high velocity movement post-stress (Schuler et al., 2025). In our data set, we examined locomotor measures during SI and observed findings consistent with these reports (Fig 4). We observed stress-related changes in movement organization in females despite the relative stability of SI ratio post-ASDS (Fig 4B,C). Specifically, we found a robust decrease in speed and acceleration between SI sessions (Fig S4). We calculated this change in acceleration and speed (no target trail speed – target trial speed = delta speed Δ) for all mice and observed a lower delta value for females following ASDS compared to controls indicating less change in speed with the target present. This maintenance of high speed may be consistent with darting or other high velocity stress-related movement within the SI arena when the target is present and females feel threatened. Indeed, there is evidence for overlap between fear and stress based behavioral and neural signatures (Ayash et al., 2023; Daviu et al., 2019). However post-CSDS we did not observe the same differences (Fig 4G-H). These findings suggest that females display immediate locomotor reactivity following acute stress, but that this response does not persist after chronic stress. Thus, the female-specific behavioral signature appears to characterize the initial response to stress rather than a stable long-term phenotype. Sex-typical behaviors, which reflect differences in natural behavioral repertoires, may influence how males and females respond to the same environmental challenge. This may be particularly relevant to our finding that females displayed more female-typical behaviors following acute stress in the form of distinct locomotion profiles (Fig 4). The divergent response to acute stress may reflect differences in how males and females perceive and respond to the social threat. Exposure to an aggressive conspecific is an environmental experience that is more natural for male mice but an unexpected or atypical threat for female mice and therefore the initial response to this acute social stressor may be shaped by sex-specific innate behavioral strategies or threat responses. With repeated exposure to the stressor, however, these initial responses may change, potentially resulting in a greater expression of behaviors typically observed in males.

The findings presented in this study are also consistent with findings of sex differences in response to other stressors. In a learned helpless model of depression, females do not display helplessness to the same degree as males (Dalla et al., 2008). Similarly, foot shocks produced divergent defensive behaviors in males and females in addition to differential patterns of c-Fos activation during social approach following the foot-shocks (Dorofeikova et al., 2023). In rats, sex differences been reported in risk mitigation strategies in response to threat of foot shock (Pellman et al., 2017). Relatedly, unlike males, females in a fear-safety-reward discrimination task did not suppress conditioned freezing following footstock in the presence of safety cue, but did decrease the number of darts (Greiner et al., 2019). Similar differences in motivation behavior have been reported following social stress where females decrease operant responding while males displayed an increase in the same behavior (McGraw et al., 2025). Divergence in the manifestation and response to stress between sexes are not only limited to adulthood but have also been reported for early life and adolescent stress (Santos-Silva et al., 2024). Females displayed increased susceptibility to depressive-like behavior following early-life stress and sex-specific changes in frontal-limbic connectivity (Goodwill et al., 2019). Differences in circulating sex hormones have often been cited as a potential driving force behind sex specific behavioral responses, however, some of the above studies carried out the same tests on ovariectomized females and determined that many of the observed differences in stress responses and strategies are independent of sex hormones (Dalla et al., 2008; Pellman et al., 2017). These works highlight key differences in post-stress behavior between males and females.

A benefit of the novel paired male-female social defeat paradigm used in the present study is that male and female mice undergo the same defeat sessions and as a result, each animal is exposed not only to direct physical aggression but also to the experience of witnessing its partner mouse being attacked, incorporating features of both social defeat and witness defeat into a single paradigm (Iñiguez et al., 2018). However, the presence of the partner mouse during defeat could also introduce additional variables and potentially buffer the effects of stress. Social buffering describes the phenomenon whereby the presence and interactions with a conspecific can have beneficial effects in the attenuation fear and stress related behaviors (Sarapultsev et al., 2026). Importantly, it been suggested that mutual experience is critical for social buffering (Parise et al., 2022). Adolescent mice exposed to physical stress showed decreased social avoidance when housed with a conspecific that had a mutual stressful experience (Parise et al., 2022). Comparably, mice that underwent contextual fear conditioning and are re-exposed to the context in the presence of an ovariectomized female cage mate display lower levels of conditioned freezing in response to the conditioned context (Gutzeit et al., 2020). In our data, the proportion of susceptibility we observed in our defeat mice post-ASDS and post-CSDS was less than has been previously observed and this could be explained by a stress buffering effect due to mutual presence of male and female subjects during defeat. In addition, CSDS has been consistently shown to induce long-term depressive like behaviors, Furthermore, CSDS has been consistently shown to induce long-term depressive like behaviors yet in our measure of depressive-like behavior or anhedonia we did not observe a significant change in sucrose preference post-CSDS (Fig S3D). Likewise, we only observed significant decrease in time spent in the open arms and marginally significant decrease in time spent in the light zone with no significant change in other anxiety-like behaviors (Fig S3A-C). Based on previous findings would have expected a greater increase in other anxiety like behaviors and this attenuation of anxiety-like and depressive like behaviors could also be explained by a buffering effect of this paired paradigm. Another strength of this study is the longitudinal design, which allowed acute and chronic stress effects to be examined within the same behavioral framework (Fig 1). The inclusion of multiple behavioral assays also provided a broader picture of affective change than social interaction testing alone. However, although the behavioral data support sex-dependent divergence, the study is currently limited to behavioral phenotyping and does not yet identify the neural mechanisms underlying these effects. Future work should aim to expand these findings to incorporate different female social stress models and parallel analysis into the underlying neural circuitry.

Overall, the data support a model in which males and females both respond to social defeat stress, but they do so in behaviorally distinct ways. In males, changes in social approach/avoidance remains a robust marker of both acute stress effects and chronic stress susceptibility. In females, stress-related changes appear to be expressed less clearly through standard SI ratio and may instead emerge through alternative behavioral dimensions including locomotor organization and broader multivariate behavioral profiles. These findings underscore the need to refine female-appropriate outcome measures in stress research and highlight the importance of characterizing early behavioral responses if the field is to move toward better prediction of vulnerability and improved prevention-focused strategies.

## Conclusion

The exclusion of female subjects from preclinical work has been a major source of bias leading to an inadequate understanding of disease manifestations in females resulting in potential mis- and underdiagnoses (Merone et al., 2022; Rechlin et al., 2022). While standard practice has shifted towards the inclusion of female subject and there has been significant efforts to modify traditional paradigms of social defeat stress to facilitate female directed aggression and defeat, the criteria and outcome measures originally defined in male cohorts has simply been extended to female subjects without revaluation or consideration for sex-specific stress response (Woitowich et al., 2020). This is particularly problematic where behavioral observations are used to classify mice and the “absence” of specific behavior or behavioral change is treated as the absence of an effect or phenotype. The data presented in this study demonstrate clear evidence of a female specific stress outcomes, highlighting that classic outcome measure such as SI ratio may not reliably capture female stress responses. This begs for a critical need to reinterpret the criteria used to classify stress responses in female mice.

## Author Contributions

DD, KA conceptualize the study. DD, KA, AM design the experiments. ES, AM, KA, AL, SJ, DF, LB, LA, PR, VS conducted the behavioral experiments. ES and AM analyzed the data and wrote the manuscript. ES, AM, DD reviewed and finalize the manuscript. DD supervised the study.

## Acknowledgements

We thank Casey Lardner for her contributions to study design and data acquisition. We thank past and present members of the Dumitriu Lab for scientific discussions and support.

## Supplementary figures

**Supplementary Figure 1:**
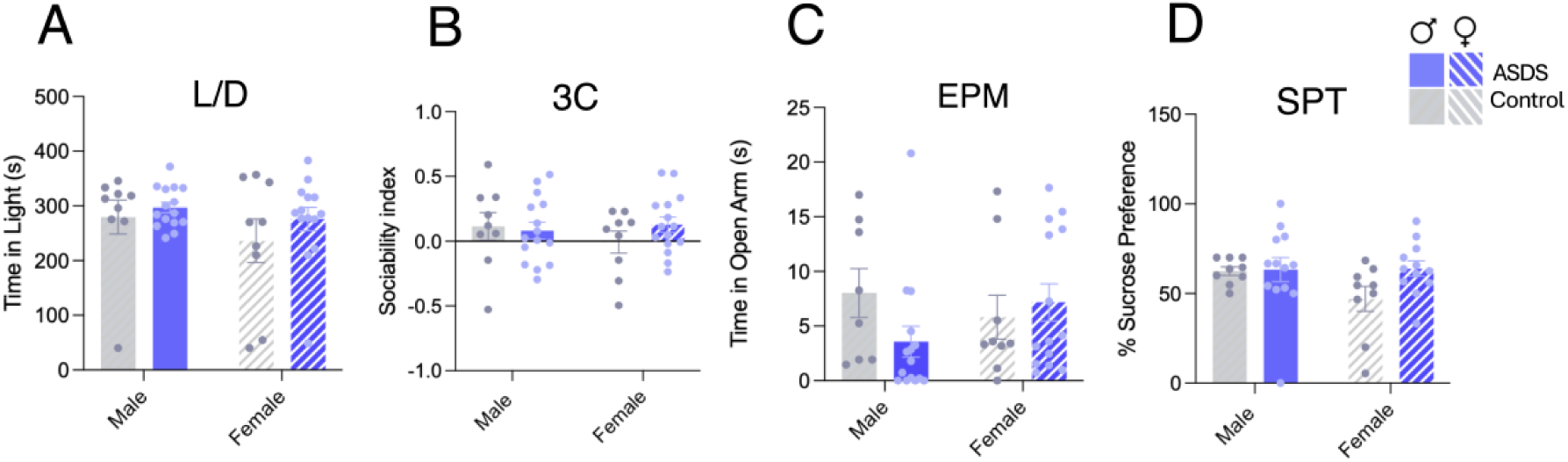
(A-D) Quantification of anxiety and depression battery measures post-ASDS (A) Quantification of raw time in light zone in control and defeated mice (B) Quantification of sociability index in control and defeat mice (C) Quantification raw time in open arms on the EPM in control and defeated mice (D) Quantification of the percent of sucrose preference in control and defeated mice. Data presented as mean +/-SEM. Statistical comparison performed using (A-D) Two-way ANOVA with Bonferroni post-hoc.

**Supplementary Figure 2:**
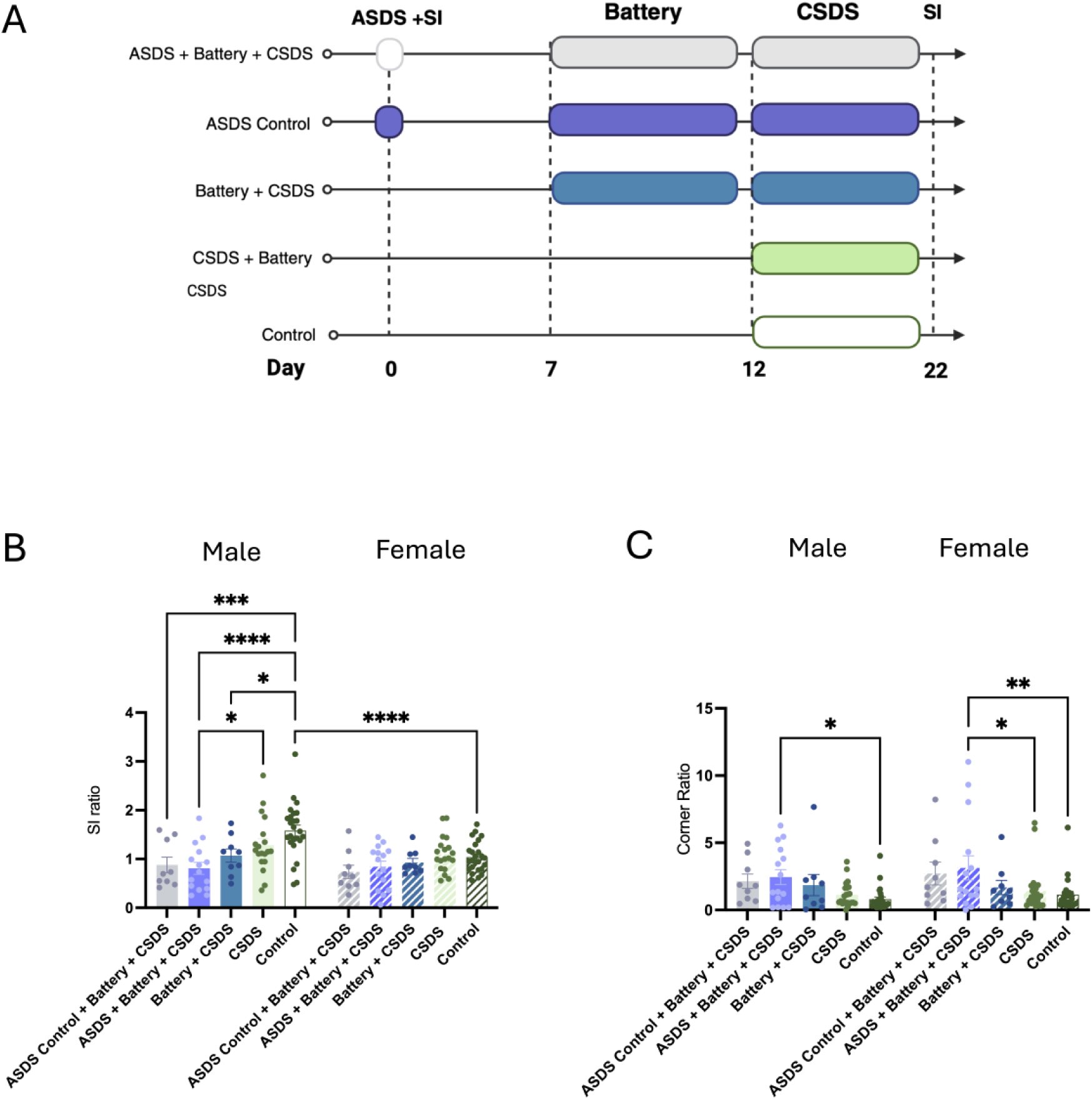
Individual group comparison of SI and corner ratio post CSDS. (A) Schematic of experimental groups and timeline. (B) Quantification of SI ratio post-CSDS. (C) Quantification of Corner ratio post-CSDS. Data are presented as mean in +/-SEM. Statistical comparison performed using Two-way ANOVA with Bonferroni post-hoc *P < 0.05, **P < 0.01, *** P < 0.001, **** P < 0.0001.

**Supplementary Figure 3:**
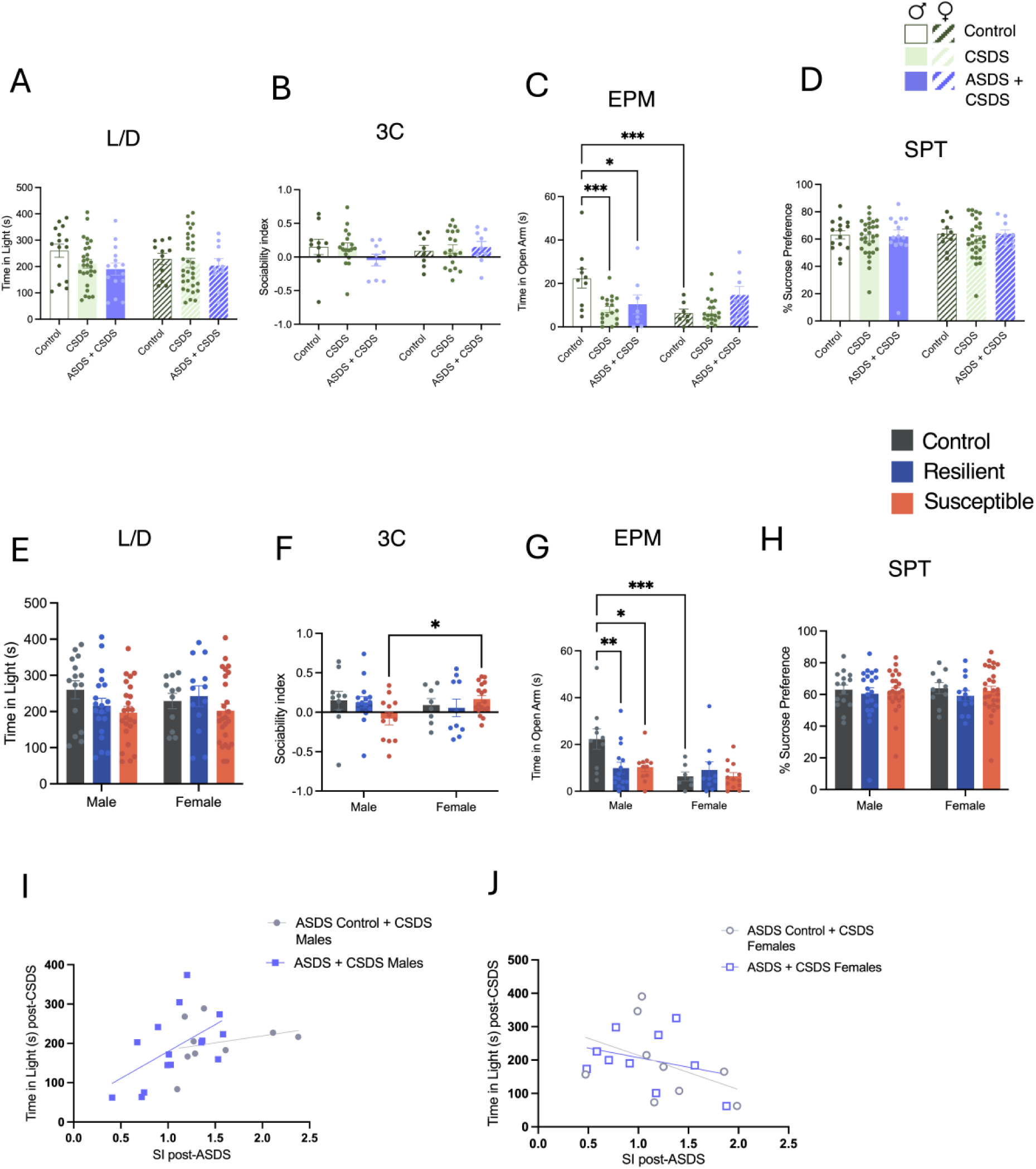
(A-D) Quantification of anxiety and depression battery measures post CSDS. (A) Quantification of raw time in light zone in control and defeated mice (B) Quantification of sociability index in control and defeat mice (C) Quantification raw time in open arms on the EPM in control and defeated mice (D) Quantification of the percent of sucrose preference in control and defeated mice. (E-H) Quantification of anxiety and depression battery measures in control, resilient and susceptible mice. (I-J) Linear regression of time in light zone in L/D test post-CSDS and SI post-ASDS. Data presented as mean +/-SEM. Statistical comparison performed using (A-H) Two-way ANOVA with Bonferroni post-hoc *P< 0.05, **P < 0.01, *** P < 0.001 (I-J) Pearson correlation *P< 0.05.

**Supplementary Figure 4:**
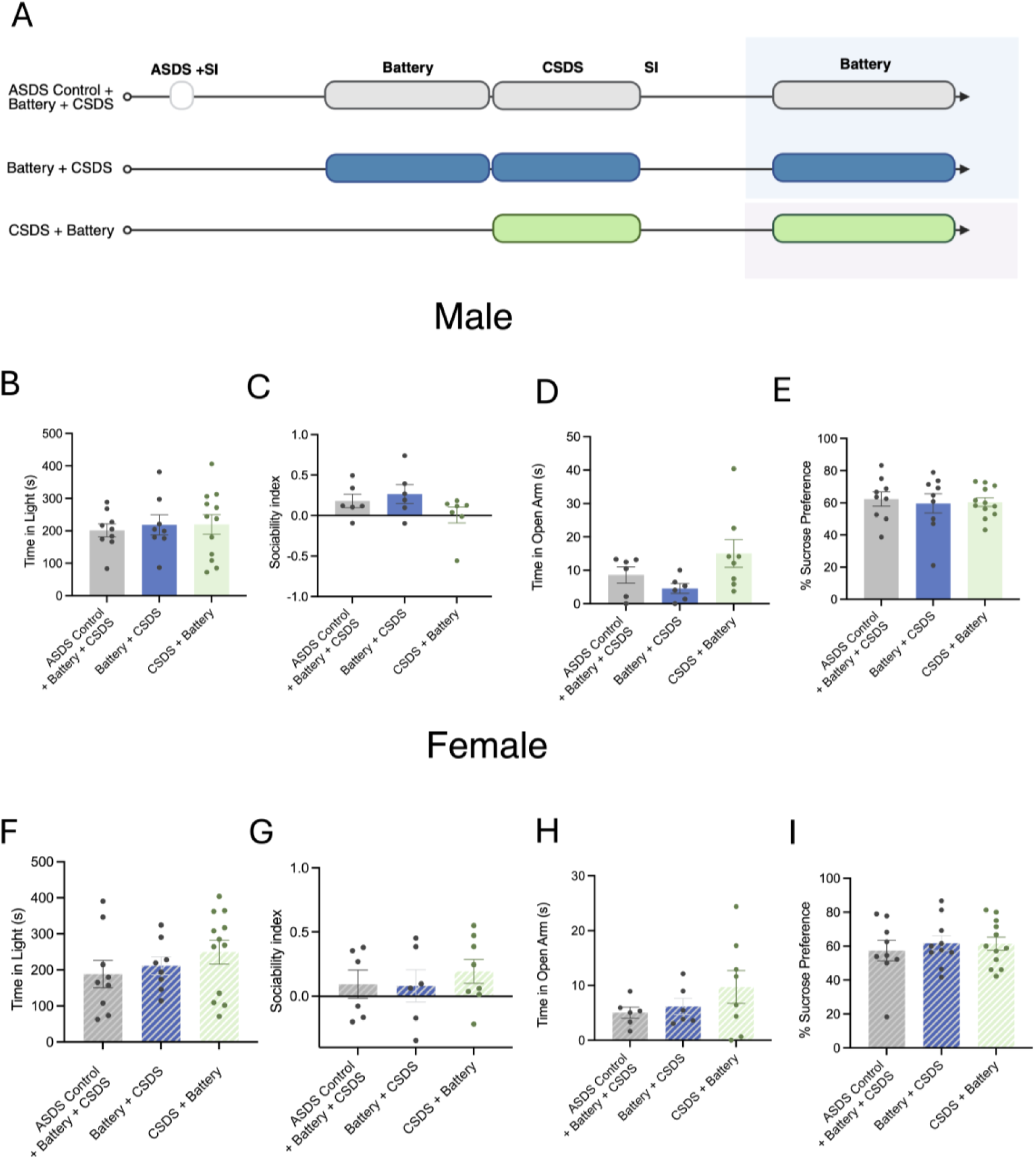
No difference in anxiety measures between Control ASDS, Battery+ CSDS and CSDS only groups. (A) Schematic of experimental groups. (B-I) Quantification of anxiety and depression measures in 3 CSDS subgroups. Data are presented as mean +/-SEM.

**Supplementary Figure 5.**
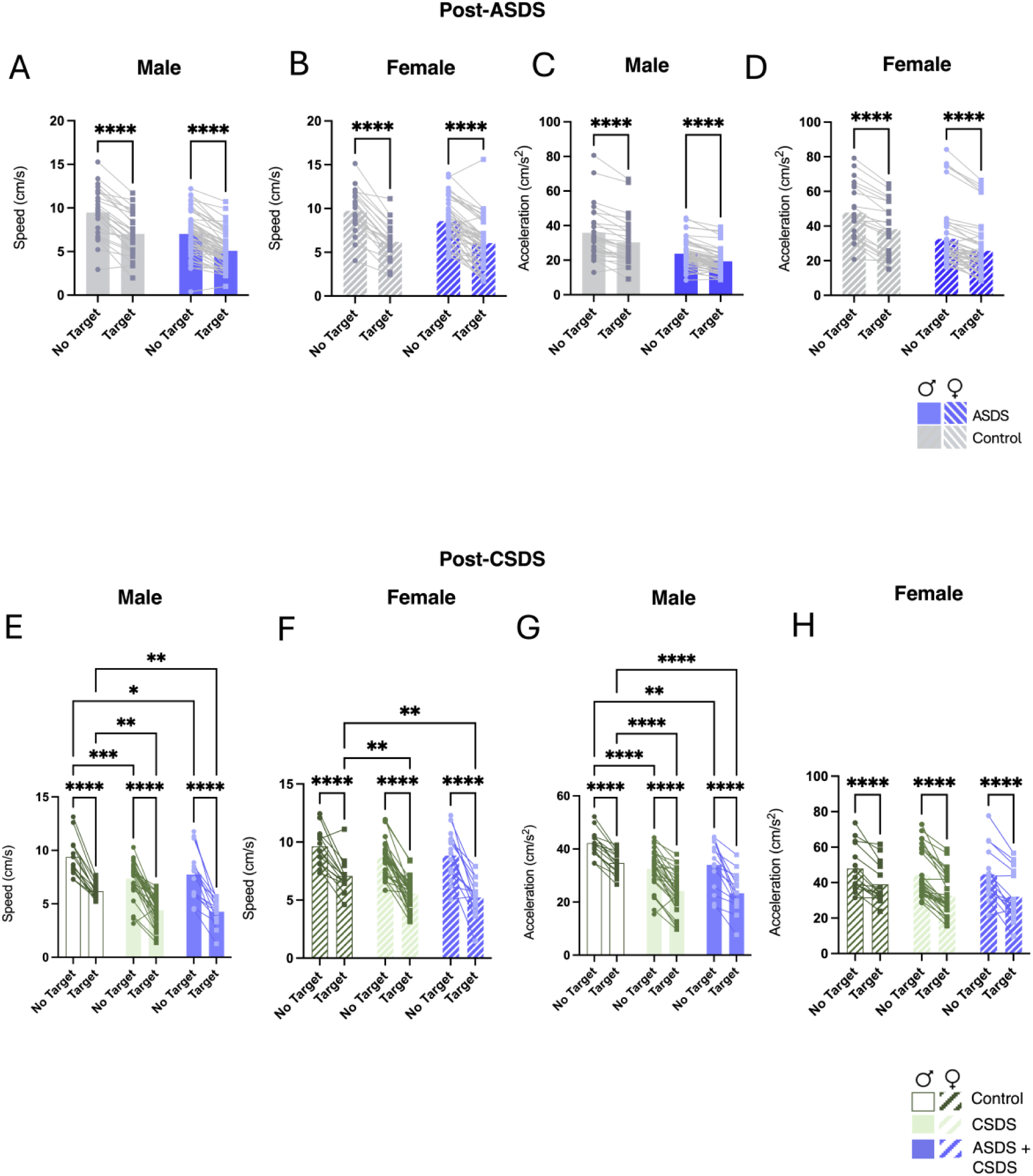
Raw Speed and acceleration across sessions following ASDS and CSDS. (A-D) Post-ASDS speed (A-B) and acceleration (C-D) measured during the no-target and target session of the SI test in male and female control and ASDS-exposed mice. (E-H) Post-CSDS speed (E-F) and acceleration (G-H) measured during the no-target and target phases of the social interaction test in male and female control, CSDS, and ASDS + CSDS groups. Data are presented as mean +/-SEM. Statistical comparison performed using RM Two-way ANOVA *P < 0.05, **P < 0.01, *** P < 0.001, **** P < 0.0001.

**Supplementary Figure 6:**
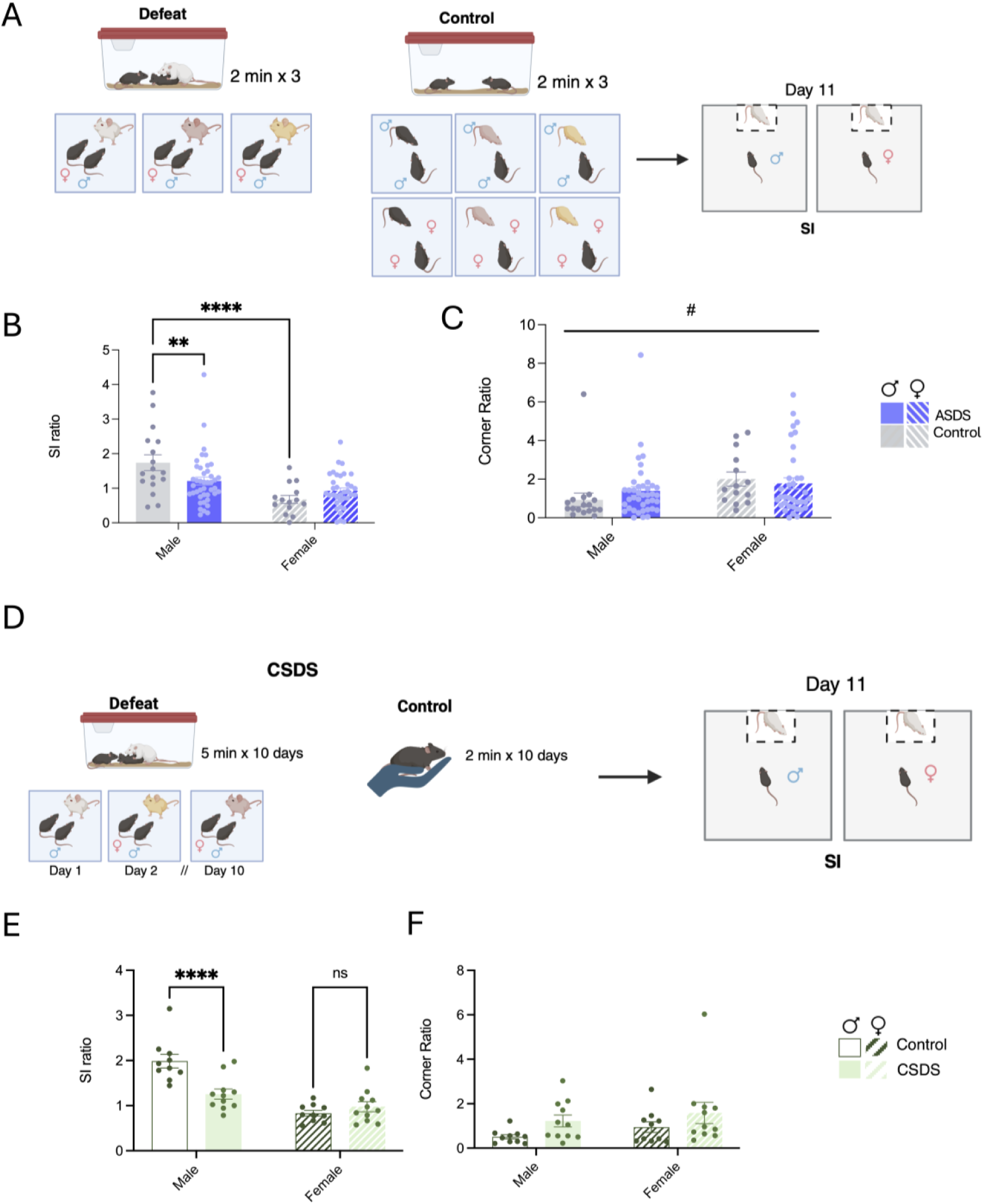
(A) Schematic of ASDS/control behavioral protocol for this specific cohort of ASDS mice. (B) Quantification of SI ratio post-ASDS. (C) Quantification of Corner ratio post-ASDS. (D) Schematic of CSDS/control behavioral protocol for this specific cohort of ASDS mice (E) Quantification of SI ratio post-CSDS. Quantification of corner ratio post-CSDS. Data are presented as mean +/-SEM. Statistical comparison performed using Two-way ANOVA Main effect: #P < 0.05. Post-hoc: *P < 0.05, **P < 0.01, *** P < 0.001, **** P < 0.0001.

